# A Repulsive Bias in Perceived Heading Away from the Straight-Ahead as Determined by Continuous Psychophysics

**DOI:** 10.64898/2026.10.07.757258

**Authors:** Mahir Rafi, Laurence R. Harris, Björn Jörges

## Abstract

Previous studies of heading perception have reported both attraction toward and repulsion away from straight ahead. Different potential explanations for these differences have been brought forward, such as the reference frame of the response (where egocentric responses are connected to attraction and allocentric responses are connected to repulsion) or the response range (where a wider response range is related to more repulsion). In this study, we aimed to contribute to this debate using a novel Continuous Psychophysics-based task in virtual reality to investigate systematic heading biases across the full 360° range (repulsion predicted) in an egocentric task (attraction predicted), while concurrently examining the perceptual and motor contributions to these biases. Participants (n = 60 across two experiments) continuously indicated their perceived heading using a joystick while experiencing optic flow consistent with slow or fast self-motion while the simulated heading direction was changing continuously. In a CONTROL condition, participants aligned the joystick with the direction of a clearly visible arrow. The joystick was placed on the dominant-hand side in Experiment 1 (on the right for n = 28 participants) and in the middle and on the left in Experiment 2. In both experiments, we found consistent repulsion away from straight ahead in the CONTROL, FAST, and SLOW conditions, regardless of joystick position. Biases around the other cardinal directions were less consistent and generally tended to show attraction. Around straight ahead, we also observed a general progression from smaller biases in CONTROL to larger biases in FAST and SLOW, suggesting that decreased reliability of perceptual cues increased biases. The broad pattern of findings remained reasonably consistent across the three joystick positions. Our original prediction of attraction in an egocentric task was therefore not supported. Instead, the findings were more consistent with the response-range account, which links larger response ranges (like our 360° response range) to stronger repulsion. Our findings further support the use of Continuous Psychophysics to study systematic perceptual biases and provide a blueprint for the analytical steps required to do so.

## Introduction

Heading perception refers to the implicit and explicit ability to determine the direction in which we are moving through the environment around us. It is a multisensory process consisting of visual and nonvisual cues. Among these cues, humans are commonly thought to be most sensitive to visual cues, especially optic flow (Cuturi & MacNeilage, 2013; Jörges et al., 2024) as well as vestibular cues (Rosenblum et al., 2022). However, recent studies have indicated that modalities such as haptic cues (Rosenblum et al., 2022) or efference copy information (Telford et al., 1995) also contribute to heading perception.

Even though our heading perception is relatively accurate in our daily life and allows us to perform a variety of tasks with great accuracy and precision, numerous studies have found systematic biases in this perceptual skill, particularly in relation to the straight-ahead. The findings of these studies are contradictory, with some studies reporting repulsion away from the straight-ahead (Crane, 2012; Cuturi & MacNeilage, 2013; Gu et al., 2010) and others reporting attraction toward the straight-ahead (D’avossa & Kersten, 1996; Sun et al., 2024; Wang et al., 2022; Warren & Kurtz, 1992; Warren & Saunders, 1995).

The reference frame of the response has emerged as a strong contender (Reisenegger et al., 2025): Repulsion away from the straight-ahead, i.e., a bias to perceive heading more lateral than the stimulation would indicate, appears to be associated with allocentric response modalities, and attraction to the straight-ahead, i.e., a bias to perceive heading closer to the straight-ahead than the true stimulus value, may be associated with egocentric tasks. Egocentric tasks require participants to respond based on visual stimuli with reference to themselves. Conversely, in allocentric tasks, participants responded with reference to the surrounding environment, e.g., by forcing a perspective change to a a bird’s-eye viewpoint. As an example of an allocentric task, Cuturi and MacNeilage (Cuturi & MacNeilage, 2013) investigated heading perception biases using visual and/or vestibular stimulus. Participants performed a heading discrimination task while seated on a motion platform simulating linear motion and/or receiving visual stimulation and reported their perceived heading (allocentrically) using a 360° arrow adjustment method The results of this study consistently showed a repulsive bias away from straight-ahead for visual, vestibular and visuo-vestibular conditions. A paradigmatic example of an egocentric task can be found in a more recent study by Sun et al. (2023): participants were moved through a starfield and then had to indicate their perceived heading (egocentrically) by moving a blue vertical line along a horizontal line projected in front of them. An attraction towards the straight-ahead was reported.

Another strong candidate cause for the discrepancy between repulsion and attraction has recently been brought forward by Sun et al (2025): Over a series of three experiments, the response *range* (rather than the response *method*) emerged as the most likely underlying factor; in their Experiment 3, participants responded to heading directions from the interval [-40°; 40°] using an arc of three different sizes (80°, 160°, 240°) to record responses while keeping all other factors identical. This included the range of possible stimuli, which was set to +/-40° for all three response ranges. The bias shifted from attraction (80°) to increasing repulsion (160° and 240°) as the response range increased. While this is a plausible explanation in and of itself, we would also like to point out that those response modes where Sun et al. (2025) report repulsion are either clearly egocentric (line) or can be interpreted as egocentric-leaning (80° arc, which is similar enough to the line to evoke an egocentric frame of reference), while the response modes where attraction was found (full circle, 160° and 240° arc) are better described as allocentric. On top of their response range-based explanation, their results are therefore also compatible with Reisenegger’s explanation based on egocentric versus allocentric responses.

Our study aims to add to this debate with a novel task based on Continuous Psychophysics (Bonnen et al., 2015). Our task has several distinguishing features that will contribute to settling the question of what is underlying the contradicting results in the literature: first, unlike any of the other studies in the area, we are coupling continuous stimulation with a continuous response. By adding this novel task to the body of evidence, we test and expand the limits of generalizability of earlier findings. Second, we are presenting participants with a full 360° of heading directions, rather than focusing on a more or less narrow band of heading angles around the straight-ahead. Third, we decouple a potential natural prior for straight-ahead heading directions from the experimentally induced prior, allowing us to draw conclusions as to whether biases regarding the straight-ahead are naturally occurring or due to common experimental designs.

Continuous Psychophysics (Bonnen et al., 2015) is an innovative alternative to traditional trial-based methods for data collection. In this paradigm, participants respond continuously to a continuously changing the stimulus are recorded. Among others, this can provide more flexibility for experiment design, speed up data collection, be less torturous for participants and provide a more realistic and natural assessment of perceptual processes (Bonnen et al., 2015). In their short history, Continuous Psychophysics experiments have largely been used to assess temporal aspects of perception using cross-correlation analyses as well as sensory precision using Kalman filter approaches. Studies using Continuous Psychophysics to investigate *biases* (i.e., accuracy) are, however, notably absent from the literature. A secondary goal of this study is therefore to explore the potential of this method and identify associated methodological challenges in assessing perceptuo-motor accuracy.

One potential pitfall of this approach is that motor noise constitutes a larger share of the overall variance in responses. For instance Straub & Rothkopf (2022) suggested that hand movements might not always capture perception precisely and accurately, which may make results obtained by this technique less reliable if not modelled appropriately. To tease apart perceptual and motor processes, we added a control condition in which participants continuously aligned a joystick with an arrow rather than their heading, minimizing the perceptual component of the task. To further address the potential for motor confounds, we varied the position of the joystick used to gather responses across two experiments. In the first experiment, we placed the joystick on the side of the participant’s dominant hand (i.e., mostly on their right). In the second experiment, participants performed the task with the joystick centered in front of their bodies and on their left.

In this study (comprised of two experiments), our primary goal is to investigate whether participants show systematic biases in perceived heading direction when using a Continuous Psychophysics task with a joystick. Specifically, we aim to understand whether our task, which uses an egocentric frame of reference, produces attraction biases toward the straight-ahead direction. Based on previous research showing attraction in egocentric tasks and repulsion in allocentric designs as per Reisenegger et al. (forthcoming), we hypothesized that participants in our egocentric task would show an attraction bias toward straight-ahead. Alternatively, given our 360° response range, repulsion might be expected as per Sun et al. (2025)). The motor control condition further allowed us to disentangle biases induced by the motor response from biases induced by heading perception proper.

A secondary, methodological goal was to validate the use of Continuous Psychophysics to study perceptual biases, beyond its historical use to assess temporal aspects of perception and sensory precision.

## Methods

### Participants

For Experiment 1, we recruited n=30 participants from the York University psychology undergraduate participant pool. They all received course credit as compensation. All participants had normal or corrected-to-normal vision and had no known history of vestibular disorders. 28 participants were right-handed and 2 were left-handed. For Experiment 2, data were collected between January and March 2026. An additional n = 30 participants were recruited and compensated using the same process described in Experiment 1. Data collection for this experiment occurred in December 2024 and January 2025. For Experiment 2, all the participants were right-handed, and data collection occurred between January and March 2026. The study was approved by the York University Office for Research Ethics (ORE) under the protocol numbers #e2019-447 (first experiment) and #e2025-303 (second experiment). Written consent was obtained from all participants before starting the experiment and the study was conducted in accordance with the principles of the Declaration of Helsinki.

### Apparatus

The same apparatus was used for both Experiment 1 and 2. We used an Alienware laptop with 16 GB of RAM, an Intel Core i7-9750H CPU (2.60 GHz), an NVIDIA GeForce RTX 2060, and a VIVE Pro EYE head-mounted device (which has a 110° field of vision, 1540 × 1600 resolution per eye, and a refresh rate of 90 Hz) to present the stimulus, which was programmed in Unity (2021.3.13f). The Unity project (https://osf.io/yskgd) and the executable (https://osf.io/6fr5t) are available from the Open Science Foundation. Participants used a Logitech G Extreme 3D PRO joystick to respond, with inputs recorded by Unity at 90 Hz. Data analysis was carried out in R (version 4.3.2) (R Core Team, 2017).

#### Stimulus

A virtual environment was created with a uniformly white ground plane and potted plants to provide visual features in the environment that would create a convincing sense of motion when a person was simulated as moving through the environment (see Figure 1). Participants were immersed in this environment and experienced 180 s of visual self-motion along one of the nine pre-generated paths (see Figure 2 for an example of such path). There was no simulation of rotation, and participants always faced the same direction. R was used to pre-generate the paths based on three guidelines: (1) The participants started in a random direction. (2) A variation in direction was selected at random following a random walk procedure, i.e., every 200 ms a new angle was chosen from a distribution with a mean of 0° and a standard deviation of 20° and added to the current presented heading angle. (3) If the next 200 ms of motion would have taken participants beyond a specified circular boundary, 180° was added to the chosen heading angle, causing them to continue in the exact opposite direction. This was done so that future versions of the same experiment might be carried out on a MOOG motion platform with a limited range of motion. Figure 2 shows the presented heading angles over the course of one 180 s run (A) as well as the distribution of heading angles presented throughout one such run (B). The R script used to generate these pathways is available in the project’s repository on Open Science Foundation (https://osf.io/au6ch/).

**Figure 1.**
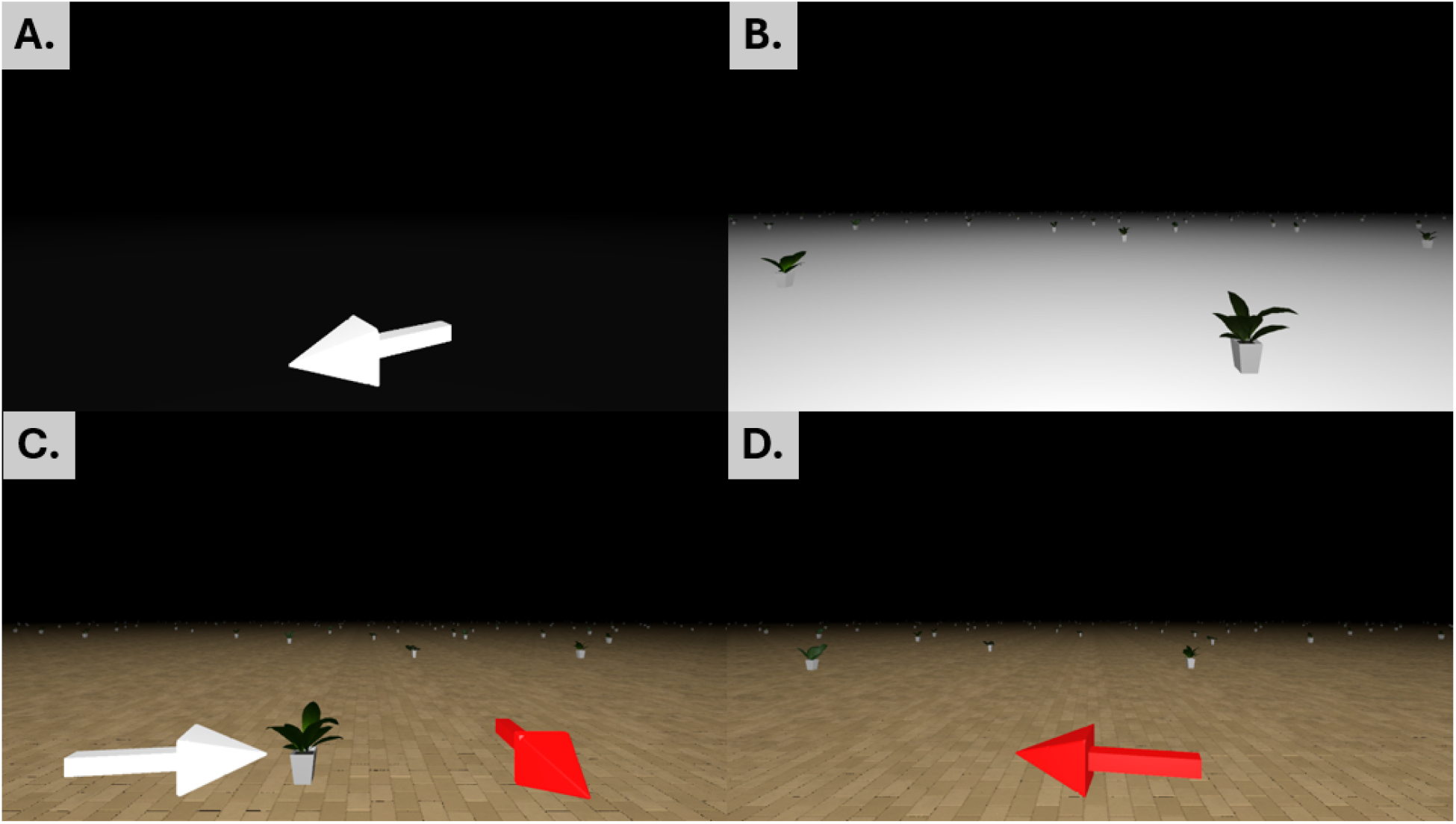
Screenshots from the experiment. A. motor control condition, B. SLOW/FAST heading condition. C. First training. D. Second training.

**Figure 2.**
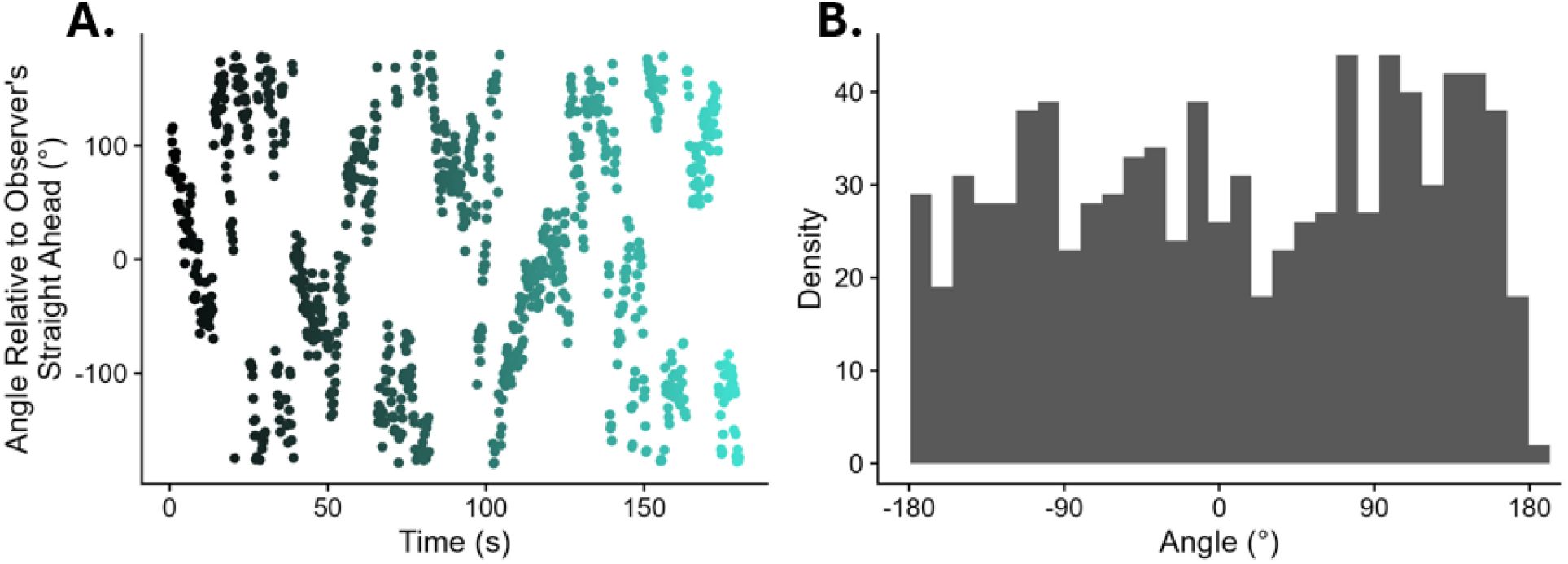
A.Heading angle presented to the observer over one run. The time in the run is on the x axis and color-coded, and the angle (relative to the observer’s straight-ahead is on the y axis. B. Distribution of presented angles across one run.

In our study, participants experienced optic flow consistent with varying self-motion speeds (SLOW and FAST) and they were asked to align a joystick (positioned where they could comfortably reach it with their dominant hand; for 28 participants it was on the right side and for 2 participants it was on the left side; see Figure 3A and B) with their heading direction. In the SLOW condition, speeds ranged from 0.4–0.6 m/s, while they ranged from 2–3 m/s in the FAST condition. In both conditions, the speed changed every 200ms between values randomly drawn from these intervals. In addition to these optic flow conditions, we introduced a motor control condition (CONTROL) where participants performed a similar task but without any optic flow. Rather, they had to align the joystick with an arrow whose direction was generated the same way as the heading stimulation in the main experimental conditions. To make all three conditions more realistic, eye height was set to 1.3 m, which is roughly the average sitting eye height, and plant size was simulated at 0.6 m. The overall design was intended to approximate a real-world experience in relevant aspects. Both the SLOW and FAST optic flow conditions included plants displayed on a flat white ground with a black background (see Figure 1B). For these conditions, 1,000 plants were randomly placed within a 10,000 m^2^ area, with a minimum spacing of 5 m between each instance to avoid clustering. The arrow in the CONTROL condition was presented in a uniformly black environment. You can download both the executable and the Unity project used to create the executable on Open Science Foundation (https://osf.io/au6ch/).

**Figure 3.**
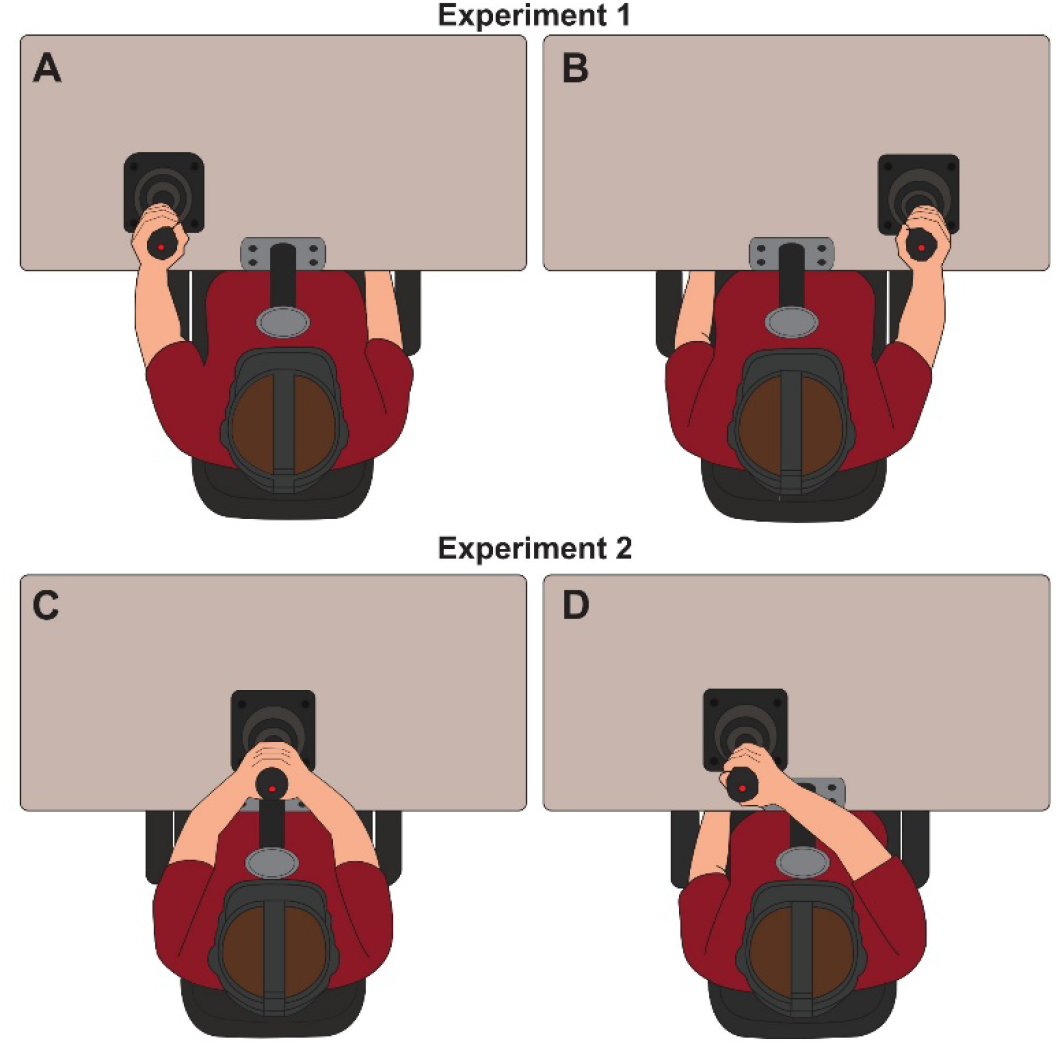
Joystick positions and hand configurations used for Experiment 1 and 2 A. Experiment 1: joystick positioned on the left side for left-handed participants, allowing them to use their dominant hand. B. Experiment 1: joystick positioned on the right side for right-handed participants, allowing them to use their dominant hand. C. Experiment 2: joystick positioned in the middle and operated with both hands. D. Experiment 2: joystick positioned on the left and operated with the dominant right hand which required participants to reach across the body.

#### Procedure

For Experiment 1, we had 5 conditions in total: 2 training conditions, 1 motor control condition and 2 heading conditions with different speeds (FAST and SLOW). Every participant started off with the training conditions. In the first training condition, participants saw two arrows: a red arrow showing the actual direction they were moving and a white arrow showing the direction they were pointing the joystick as they were getting familiarized with the task (Figure 1C). In the second training condition, the white arrow was removed and only the red arrow remained, which indicated the direction in which participants pointed the joystick based on their perceived heading perception (Figure 1D). For training, we used 1,000 plants spaced around 5 m apart with self-motion speeds ranging from 1 m/s to 1.5 m/s. To make self-motion easier to perceive, we added a tiled ground texture to the floor (Figure 1). This texture was only used in the training conditions and later removed from all other conditions to avoid ceiling effects in heading perception performance that could obscure the impact of our manipulations. There were no formal criteria for passing the training and all participants proceeded to the main experiment after completing both training conditions. After training, all participants completed one motor condition (Figure 1A) and two optic flow conditions (SLOW vs. FAST; Figure 1B), presented in random order. Overall, the study took no more than 30 minutes to complete.

**Experiment 2** – In Experiment 1, we had our joystick placed comfortably on the side of the participant’s dominant hand. 28 of them were right-handed, that is, our dataset was heavily skewed towards a right-side joystick position. Further, our setup might not be interpreted as fully egocentric: when the joystick is placed on the side of the participants body, a transformation (albeit minor) has to occur between the participant’s viewpoint (as reference for their perceived heading) and the position of the joystick (as reference for their response). To address these two issues, we conducted a second experiment, in which the joystick was either placed centered in front of the observer or on their left. The middle condition minimized the need for spatial transformations, maximizing its egocentric nature. The left condition allowed us to explore whether our results generalized to a wider range of joystick positions and served as a symmetrical counterpart to the (mostly) right-side placement of the joystick in Experiment 1. All was kept equal relative to Experiment 1, except for the following changes: In order to allocate the increased number of runs (2 trainings, 3 runs with the joystick in the middle, 3 runs with the joystick on the run), we decreased the duration of each run to 120 s. As in Experiment 1, participants first completed Training 1 and Training 2. After that, they completed three runs with the joystick placed in one position (middle or left) and three runs with the joystick placed in the other position. For the middle position, participants were asked to put both hands on the controller, and for the ition, we asked participants to use their dominant hand. Since all participants in this cohort were right-handed, this entailed reaching across their body to the joystick on the left, see Figure 3C-D. We counterbalanced whether participants started with the joystick in the middle or on the left, and within each joystick position, the order of conditions (SLOW, FAST, and CONTROL) was randomized.

### Data Analysis

To examine how participants followed the presented heading in VR with the joystick, their responses were recorded continuously while the direction of the stimulus changed every 200ms. An example from one participant is shown in Figure 4A, where the black trace shows the stimulus and the teal trace shows the responses.

**Figure 4.**
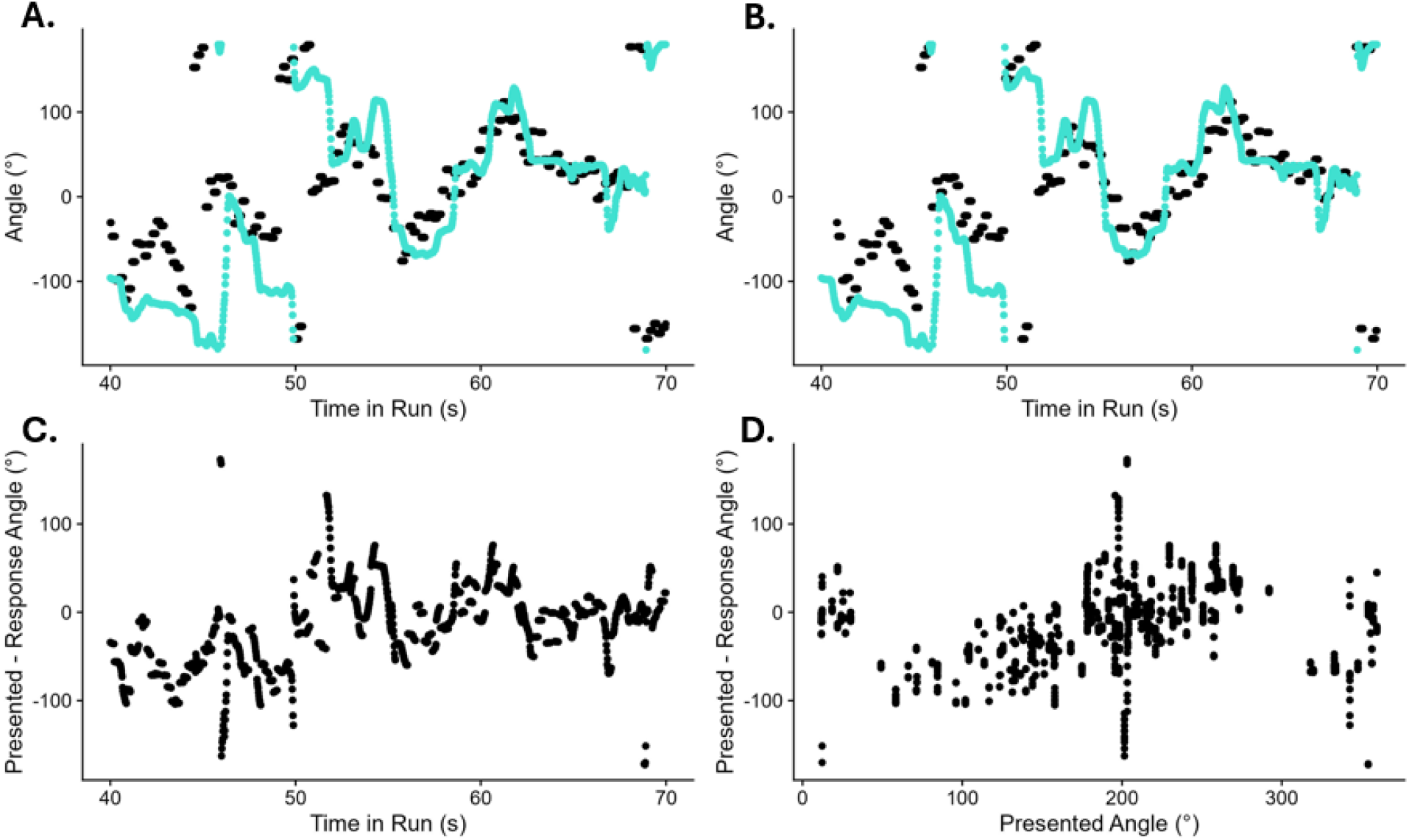
A.Stimulus and response from one participant. Black dots are the presented headings, and turquoise dots are the joystick responses B. As B, but lagged to the time lag of maximum correlation. C. Signed difference between the presented angle and the participant response angle for the time lag of maximum correlation over the course of the run D. As C, but ordered by the presented heading angles.

Because the responses were delayed compared to the stimulus, the first step was to determine the optimal time lag between the two using a cross-correlation analysis. Since heading angles are a circular variable, we first transformed both stimulus and response angles by taking their sine. Separately for each participant and condition, we then systematically shifted the responses relative to the stimulus to obtain that lag where the correlation between the two was the highest. The point with the highest correlation was taken as the optimal time lag for that participant and condition. Figure 4B shows presented angles along with the participant responses after shifting them by the optimal time lag determined for this participant and condition.

After finding the optimal time lag, we calculated the heading error as the difference between the response angle and the presented angle. Angles were coded so that straight-ahead was 0°, right was +90°, left was ™90° and back was -/+180°. Figure 4C shows the signed errors for one participant, with each point representing a single response frame against the presented angle. As next step, we ordered these same data points by the presented heading angle rather than by time in the run (Figure 4D).

Finally, to investigate whether the responses showed any systematic bias depending on the direction of the stimulus, we divided the full 360° range into four quadrants around the cardinal directions: straight-ahead (™45° to +45°), right (+45° to +135°), left (™135° to ™45°) and back (+135° to ™135°). We then examined the slope of the errors to see whether responses showed an attraction toward the center of each quadrant or a repulsion away from it. Positive slopes indicated repulsion away from the reference (straight-ahead, left, right, backwards) and negative slopes indicated attraction towards the reference.

For statistical analysis, we used Linear Mixed Modelling as implemented in the lme4 package (Bates et al., 2015) for R. The signed error (presented angle - response angle) at the optimal time lag was used as the dependent variable. We used the presented heading angle, the experimental condition (SLOW, FAST, CONTROL), as well as their interaction as fixed effects. We also included random intercepts and random slopes for the presented heading angle, the experimental condition and their interaction per participant as random effects. In Wilkinson & Rogers notation (Wilkinson & Rogers, 1973), this model read as follows:

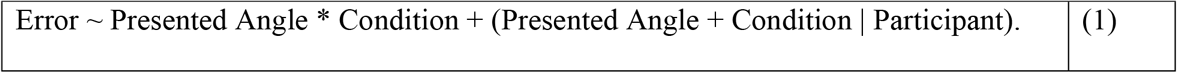

## Results

We report the results separately for each experiment/joystick position. Figure 5 visualizes biases for the full range of heading errors. This section summarizes all relevant comparisons verbally, and readers can refer to Table A1, Table A2, and Table A3 in the Appendix for all regression coefficients and 95% confidence intervals. Specifically, we report whether slopes for each condition were significantly different from zero, and whether slopes differed between conditions. The regression coefficients for CONTROL, FAST, and SLOW correspond to the slopes of the fitted lines, while coefficients comparing conditions (e.g., FAST vs CONTROL) correspond to differences between these slopes. Positive slopes indicate repulsion from the reference cardinal, while negative slopes indicate attraction towards it.

**Figure 5.**
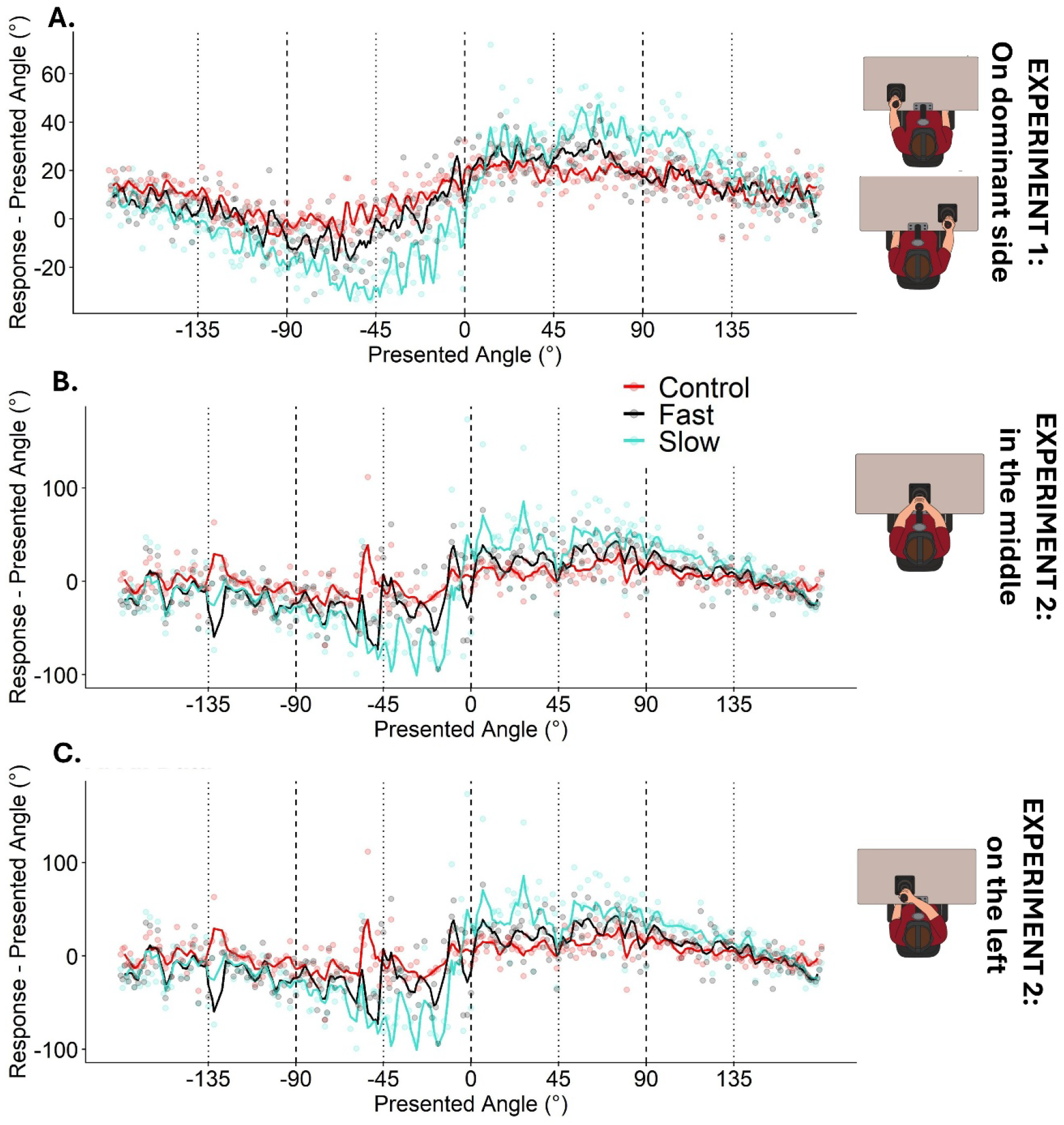
Heading errors across the full 360° range of presented heading directions. The y-axis shows the difference between the participant’s joystick response and the presented heading angle (Response – Presented Angle) at the appropriate time lag that accounts for tracking delay. Dots represent binned average errors (1° bin width), and solid lines show Gaussian-smoothed trends for each condition, both of which are color-coded (red for CONTROL, black for FAST and blue for SLOW) Vertical dashed lines show the four cardinal directions (straight-ahead, right, backward, and left) and the vertical dotted lines indicate the cut-offs between the cardinal quadrants used for statistical analysis. A. Experiment 1. B. Experiment 2: joystick in the middle. C. Experiment 2: joystick on the left.

**Experiment 1** – The results for Experiment 1 are shown in Figure 5A. For the straight-ahead, participants displayed repulsion (i.e., a positive slope significantly different from zero) for all three conditions, but no significant differences between the conditions. We found attraction for CONTROL and FAST around the right-hand cardinal, but no significant bias for SLOW, nor any differences between the three conditions. In the left-hand quadrant, FAST and SLOW were associated with attraction (i.e., negative slopes significantly different from zero), while CONTROL did not elicit a significant bias. Accordingly, we found that FAST and SLOW caused significantly more attraction than CONTROL, but they were not different from each other. Lastly, we found no significant biases around the backwards cardinal, nor any differences between the conditions.

**Experiment 2: Middle** – Figure 5B visualizes the data for those runs of Experiment 2 where the joystick was placed in the middle. For the straight-ahead, participants displayed positive slopes (i.e., repulsive biases) across all three conditions. They were stronger for FAST than for CONTROL, and even stronger for SLOW. Negative slopes (i.e., attractive biases) were found around the right for all three conditions, with no significant differences between them. Similarly, participants displayed attraction around the left, but here we did find significant differences between the conditions: SLOW was associated with stronger attraction than CONTROL and FAST, while CONTROL and FAST were not significantly different from another. For backwards, no significant bias was detected in the CONTROL condition, while both FAST and SLOW elicited attraction. Attraction was stronger for FAST than for CONTROL, as well as stronger for SLOW than for FAST.

**Experiment 2: Left** – Figure 5C visualizes the data for those runs in Experiment 2 where the joystick was placed in the middle. When the joystick was position on the participants left (Figure 5C), the found significant repulsion around the straight-ahead for all three conditions. This repulsion was stronger for FAST than for CONTROL, and significantly stronger even for SLOW. No significant biases were found around the right, while the left side elicited attraction for all three conditions. This attraction was stronger for SLOW than for CONTROL. In the backwards quadrant, CONTROL was not associated with any bias. SLOW and FAST, however, did cause attraction, and more strongly so than CONTROL.

## Discussion

In our study comprised of two experiments, we used a continuous psychophysics task in VR to investigate systematic biases in heading perception across the full 360° range. As a secondary goal, we were interested in finding out whether these biases were influenced by heading perception proper or whether they resulted from the motor response, as well as whether joystick positioning mattered. In both experiments, we found consistent repulsion away from straight-ahead for all visual conditions and different joystick positions (Figure 6). In the other cardinal directions, including leftward, rightward, and backward directions, findings were less consistent, but tended to show attraction towards the reference. Overall, these findings did not support our original hypothesis, which predicted attraction toward straight ahead based on the egocentric frame of reference of our task. Instead, these findings are more consistent with the response-range account (Sun et al., 2025), which links stronger repulsion away from the straight-ahead with larger response ranges. Interestingly, we found that biases were smaller in CONTROL, larger in FAST, and largest in SLOW (see Figure 6). This pattern suggests that perceptually more demanding conditions elicited stronger biases. Finally, the similarity of the overall findings across different joystick positions suggests that these effects were not affected strongly by the joystick placement. This also supports the use of Continuous Psychophysics as a reliable method to study perceptual biases, while at the same time providing a blueprint for the statistical procedures specific to this approach.

**Figure 6.**
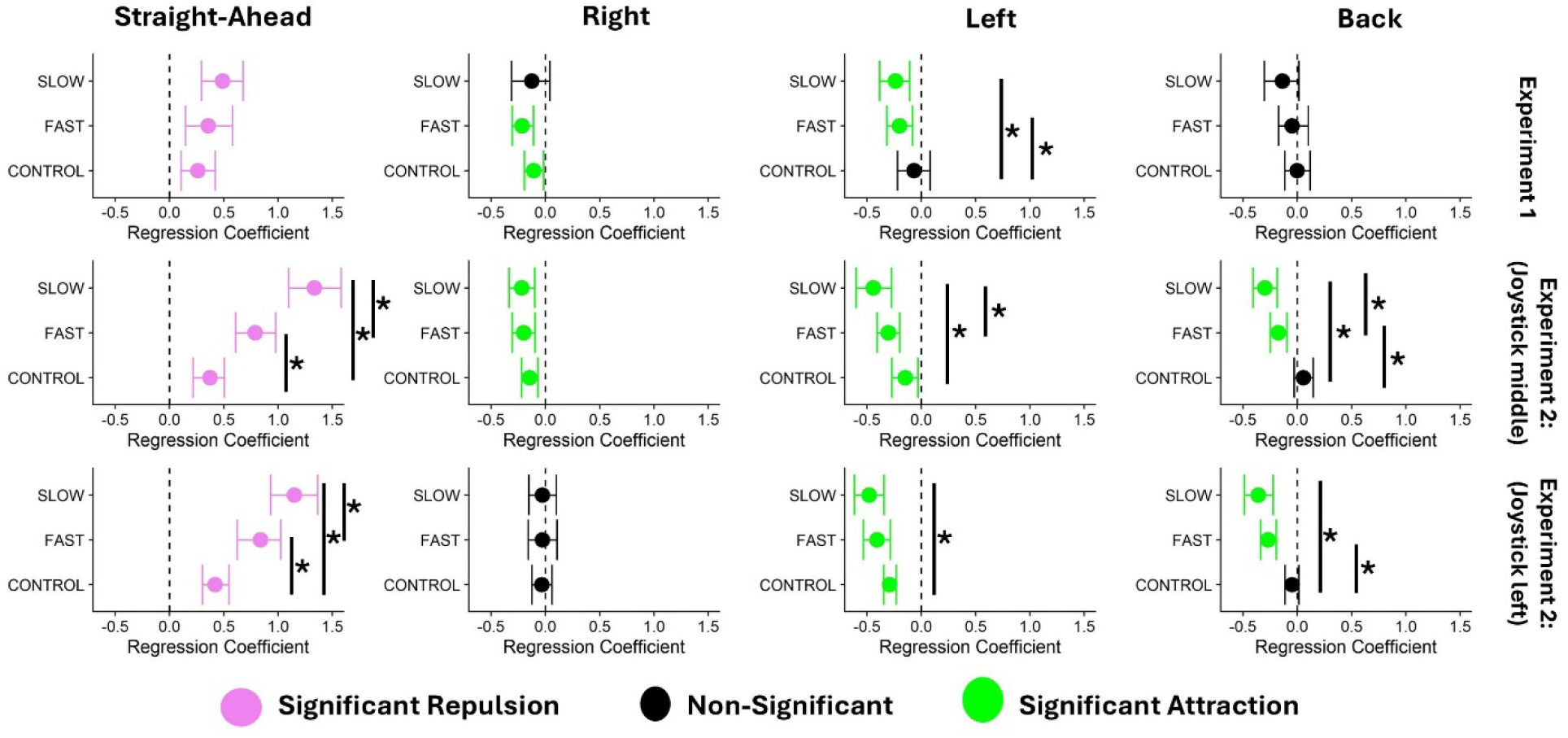
Regression coefficients (bold dots) and 95% confidence intervals (antennae) for the slopes observed in the three conditions, separately for each cardinal direction (rows of panels) and experiments and joystick positions (columns of panels). Black dots and antennae mean that the bias was not significant different from zero, pink codes for a significantly repulsion away from the reference, and green indicates a significant attraction. Significant differences between the slopes for the conditions are indicated by stars on the right of each panel. The vertical dashed line indicates a zero effect, i.e., no bias.

Our hypothesis was that participants would show attraction toward straight ahead because the task used an egocentric frame of reference, and attraction toward straight ahead had previously been reported in egocentric heading tasks (Reisenegger et al., 2025). We, however, found consistent repulsion away from the straight-ahead. This is more in line with Sun et al.’s response range account (Sun et al., 2025), which predicts repulsion for larger response ranges (like our 360° response range).

Given the characteristics of our stimulus, it is worth discussing the model brought forward recently by Sun et al. (2025) in more detail. Their contention – based on Wei and Stocker’s efficient Bayesian observer model (Wei & Stocker, 2015, 2017) – is that results in the literature regarding biases in heading perception can be explained by a push-and-pull between Bayesian processes, where responses are attracted to the straight-ahead via a Bayesian prior, and efficient coding, where efficient resource allotment around the straight-ahead would lead to a repulsion away from this reference. This percept is then scaled by the response range to result in a behavioral response. The interesting contribution of our study in the context of this model is that our stimulus values (chosen via a random walk) create a prior that is not centered around the straight-ahead. Rather, the prior consists in the expectation that the next presented heading is most likely to be similar to the current presented heading, depending on the standard deviation chosen for the random walk. While there still likely is a natural prior for straight-ahead headings because this is the most dominant perceived heading in daily life, the presence of a conflicting (experimentally induced) prior should at least lower its effects drastically. At the same time, our joystick response allows for a full 360° response range. According to Sun et al.’s model, the weaking of the attraction-inducing prior around the straight-ahead, along with a wide response range, would be expected to elicit extremely strong repulsion with regards to the straight-ahead – as found in our dataset, particularly in Experiment 2. Equally supported by our data is the efficient Bayesian observer’s prediction that higher sensory noise should lead to larger biases (Wei & Stocker, 2017), as biases tended to be stronger for SLOW than for FAST, and stronger for FAST than for CONTROL. The findings for the other cardinal directions were less consistent and tended to show attraction. It is possible that some of the attraction around the leftward and rightward cardinal directions was influenced by the strong repulsion around straight ahead, since some heading angles within these quadrants were still relatively close to straight ahead. Overall, the results suggest nonetheless that heading biases varied across the cardinal directions, as shown in Figure 6, even though the extent to which they represented influences of the straight-ahead or other cardinals is hard to ascertain using this dataset

One important part of our study was the CONTROL condition, which we included because motor factors account for a much larger share of variability in continuous psychophysics designs than for traditional, trial-based methods. In the heading conditions (FAST and SLOW), participants first had to estimate their direction of motion from optic flow and then report that direction using the joystick. In the CONTROL condition, participants still had to perceive the direction of the presented arrow, but this perceptual judgment was much easier than estimating heading from optic flow. Therefore, it is reasonable to suggest that biases observed in CONTROL reflected the motor response required to operate the joystick to a larger degree than the FAST and SLOW conditions. Generally, we did find some biases in the CONTROL condition, albeit less pronounced than in the heading conditions. This suggests either that, indeed, there are motor-induced biases at play or that object orientation judgements are subject to very similar biasing processes as heading perception.

In several quadrants and joystick positions, we observed a general progression from smaller biases in CONTROL to larger biases in FAST and SLOW, although not every comparison was statistically significant (Figure 6). This progression may reflect differences in how much directional information was available in each condition: the arrow in CONTROL provided clear and direct information, while participants had to estimate heading from optic flow in FAST and SLOW, with the SLOW condition providing less information than FAST (Jörges et al., 2024).

Another aspect of our study was to examine whether the heading biases were influenced by the position of the joystick. In Experiment 1, the joystick was placed on the side of the participant’s dominant hand. This raised the question of whether the response method was fully egocentric, or whether the off-center joystick position introduced a partly allocentric frame of reference. Further, given that 28 out of 30 participants were right-handed, the data in Experiment 1 was mostly collected with the joystick on the right, casting some doubt on whether any biases we found were perhaps just due to the joystick position. To address these concerns, we added Experiment 2, in which the joystick was placed either in the middle (to minimize necessary spatial transforms) or on the left (as a counterpart to Experiment 1’s right joystick placement). Overall, the pattern of results was consistent across all three joystick positions, especially regarding the repulsion observed around the straight ahead (Figure 6).

A secondary, methodological goal of our study was to examine whether Continuous Psychophysics, which has traditionally served to address issues of sensory precision and temporal aspects of perception (Bonnen et al., 2015; Jörges et al., 2024; Décima et al., 2022; Tonelli et al., 2025), could be used to investigate perceptual *biases* as well. In both experiments and all three joystick positions, results were reasonably consistent, especially the repulsion observed around straight ahead (see Figure 6). This consistency provides support for the use of Continuous Psychophysics to study perceptual biases, while this manuscript provides a blueprint for how to extract information about accuracy from continuous raw data. One important caveat is that Continuous Psychophysics may increase the share of motor variability in the total variability because adjusting a joystick continuously is more demanding from a motor control point of view than pressing a button once per trial as one would in traditional designs. This makes it much more important to either to capture using a modelling approach (Straub & Rothkopf, 2022), and/or to control for it experimentally. We opted for the latter by adding a condition (CONTROL) where we minimized the perceptual difficulty of the task in order to maximize the relative contribution of motor effects. Overall, the methodological conclusion is that, in opposition to traditional designs where mouse clicks are generally mostly independent of the stimulus, Continuous Psychophysics requires us to consider potential confounds for the motor component of the task as carefully as for the perceptual component.

## Conclusions

Following Reisenegger et al. (Reisenegger et al., 2025), we predicted attraction toward the straight ahead as our task used an egocentric frame of reference. However, the results did not support this hypothesis. Instead, we found consistent repulsion away from straight ahead, in line with Sun et al.’s response range account (Sun et al., 2025), with more variable biases around the other cardinal directions. Overall, our findings show that Continuous Psychophysics can be used to study perceptual biases, provided that the motor component of the response is carefully considered.

## Open Science

All data and code (both for analysis and stimulus presentation) can be downloaded from Open Science Framework (https://osf.io/au6ch/).

## Appendix

Detailed Regression Coefficients and Confidence Intervals

**Table A1.**
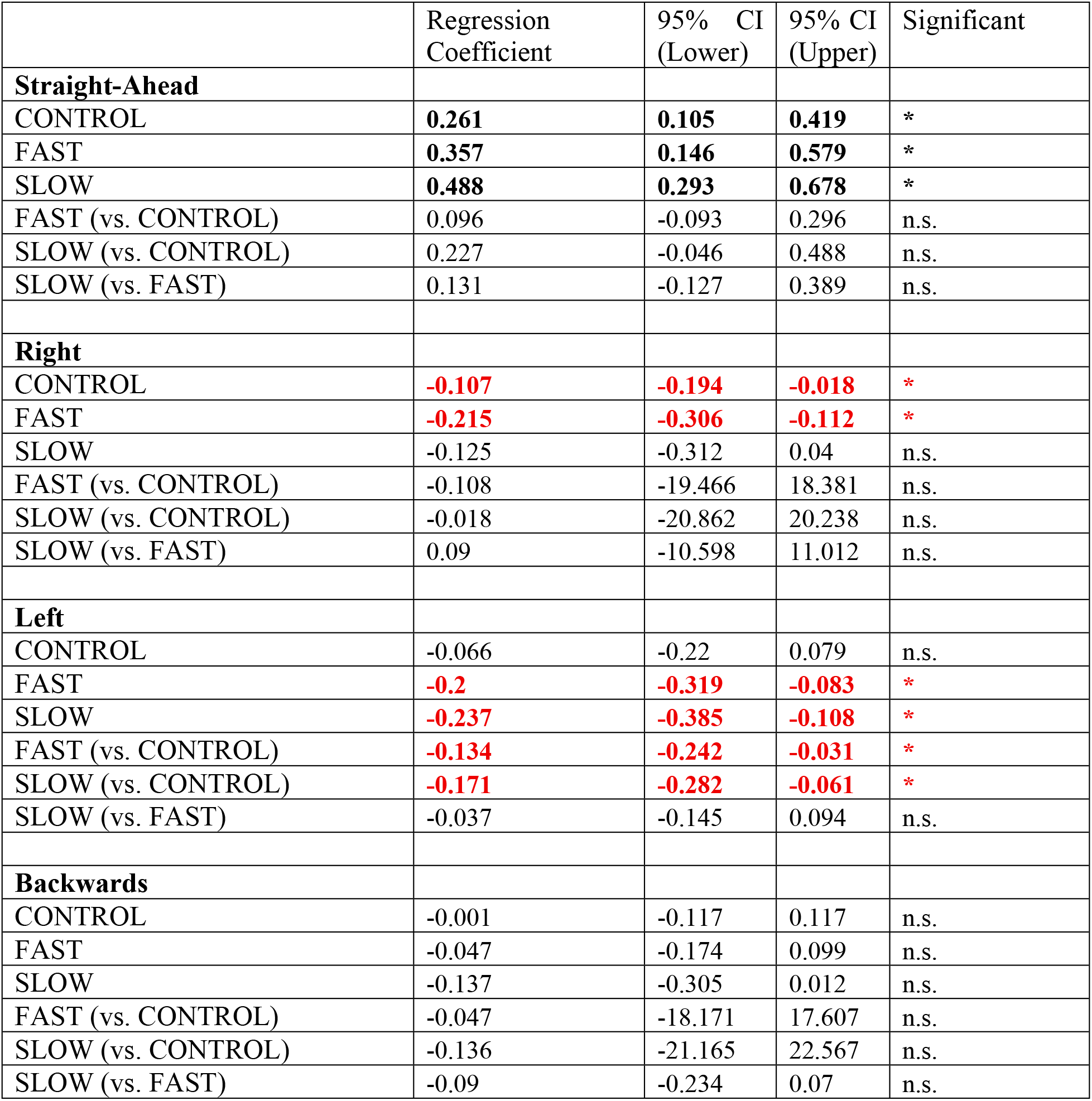
Experiment 1 (Joystick on the side of the dominant hand). Regression coefficients and 95% confidence intervals for Experiment 1, where the joystick was placed on the side of the participants’ dominant hand.

**Table A2.**
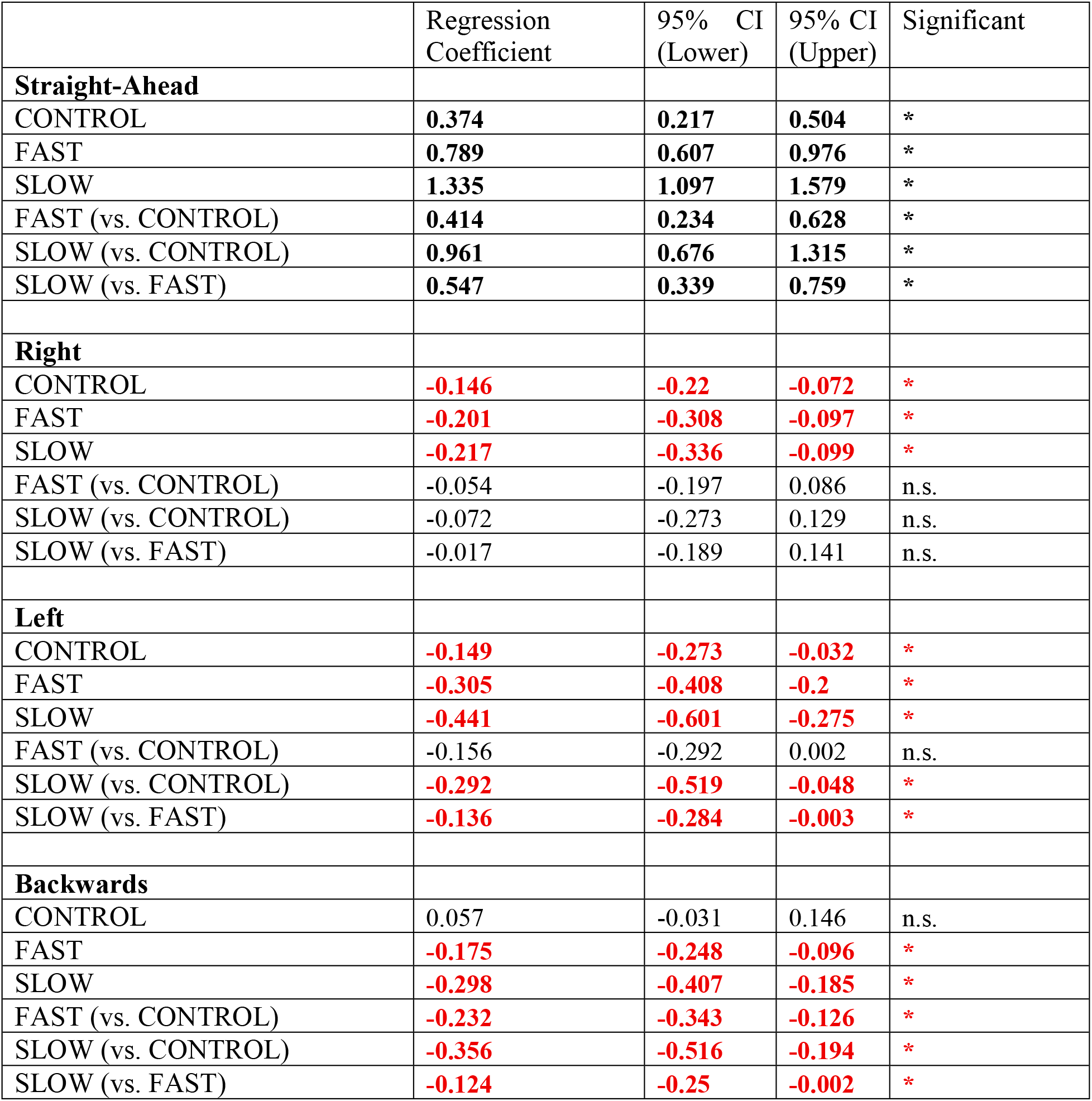
Experiment 2 (Joystick in the middle). Regression coefficients and 95% confidence intervals for those runs of Experiment 2 where the joystick was placed centered in front of the participant.

|  | Regression Coefficient | 95% CI (Lower) | 95% CI (Upper) | Significant |
| --- | --- | --- | --- | --- |
| **Straight-Ahead** |  |  |  |  |
| CONTROL | **0.374** | **0.217** | **0.504** | * |
| FAST | **0.789** | **0.607** | **0.976** | * |
| SLOW | **1.335** | **1.097** | **1.579** | * |
| FAST (vs. CONTROL) | **0.414** | **0.234** | **0.628** | * |
| SLOW (vs. CONTROL) | **0.961** | **0.676** | **1.315** | * |
| SLOW (vs. FAST) | **0.547** | **0.339** | **0.759** | * |
| **Right** |  |  |  |  |
| CONTROL | **-0.146** | **-0.22** | **-0.072** | * |
| FAST | **-0.201** | **-0.308** | **-0.097** | * |
| SLOW | **-0.217** | **-0.336** | **-0.099** | * |
| FAST (vs. CONTROL) | -0.054 | -0.197 | 0.086 | n.s. |
| SLOW (vs. CONTROL) | -0.072 | -0.273 | 0.129 | n.s. |
| SLOW (vs. FAST) | -0.017 | -0.189 | 0.141 | n.s. |
| **Left** |  |  |  |  |
| CONTROL | **-0.149** | **-0.273** | **-0.032** | * |
| FAST | **-0.305** | **-0.408** | **-0.2** | * |
| SLOW | **-0.441** | **-0.601** | **-0.275** | * |
| FAST (vs. CONTROL) | -0.156 | -0.292 | 0.002 | n.s. |
| SLOW (vs. CONTROL) | **-0.292** | **-0.519** | **-0.048** | * |
| SLOW (vs. FAST) | **-0.136** | **-0.284** | **-0.003** | * |
| **Backwards** |  |  |  |  |
| CONTROL | 0.057 | -0.031 | 0.146 | n.s. |
| FAST | **-0.175** | **-0.248** | **-0.096** | * |
| SLOW | **-0.298** | **-0.407** | **-0.185** | * |
| FAST (vs. CONTROL) | **-0.232** | **-0.343** | **-0.126** | * |
| SLOW (vs. CONTROL) | **-0.356** | **-0.516** | **-0.194** | * |
| SLOW (vs. FAST) | **-0.124** | **-0.25** | **-0.002** | * |

**Table A3.** Experiment 2 (Joystick on the left). Regression coefficients and 95% confidence intervals for those runs of Experiment 2 where the joystick was placed centered in front of the participant.

|  | Regression Coefficient | 95% CI (Lower) | 95% CI (Upper) | Significant |
| --- | --- | --- | --- | --- |
| **Straight-Ahead** |  |  |  |  |
| CONTROL | **0.421** | **0.303** | **0.547** | * |
| FAST | **0.839** | **0.623** | **1.023** | * |
| SLOW | **1.15** | **0.93** | **1.364** | * |
| FAST (vs. CONTROL) | **0.418** | **0.211** | **0.61** | * |
| SLOW (vs. CONTROL) | **0.729** | **0.504** | **0.955** | * |
| SLOW (vs. FAST) | **0.312** | **0.099** | **0.554** | * |
| **Right** |  |  |  |  |
| CONTROL | -0.033 | -0.126 | 0.06 | n.s. |
| FAST | -0.026 | -0.159 | 0.106 | n.s. |
| SLOW | -0.028 | -0.155 | 0.101 | n.s. |
| FAST (vs. CONTROL) | 0.007 | -0.133 | 0.133 | n.s. |
| SLOW (vs. CONTROL) | 0.005 | -0.179 | 0.167 | n.s. |
| SLOW (vs. FAST) | -0.002 | -0.133 | 0.137 | n.s. |
| **Left** |  |  |  |  |
| CONTROL | **-0.293** | **-0.348** | **-0.231** | * |
| FAST | **-0.409** | **-0.535** | **-0.286** | * |
| SLOW | **-0.479** | **-0.618** | **-0.344** | * |
| FAST (vs. CONTROL) | -0.116 | -0.245 | 0.016 | n.s. |
| SLOW (vs. CONTROL) | **-0.186** | **-0.314** | **-0.039** | * |
| SLOW (vs. FAST) | -0.07 | -0.183 | 0.047 | n.s. |
| **Backwards** |  |  |  |  |
| CONTROL | -0.048 | -0.113 | 0.014 | n.s. |
| FAST | **-0.269** | **-0.339** | **-0.195** | * |
| SLOW | **-0.359** | **-0.489** | **-0.223** | * |
| FAST (vs. CONTROL) | **-0.22** | **-0.332** | **-0.129** | * |
| SLOW (vs. CONTROL) | **-0.31** | **-0.464** | **-0.163** | * |
| SLOW (vs. FAST) | -0.09 | -0.206 | 0.031 | n.s. |

